# Translating antibiotic prescribing into antimicrobial resistance in the environment: a hazard characterisation case study

**DOI:** 10.1101/539536

**Authors:** Andrew C. Singer, Qiuying Xu, Virginie D.J. Keller

## Abstract

The global movement to combat the rise in antibiotic-resistant infections aims to reduce antibiotic use and misuse in humans and in food production. The environment receives antibiotics through a combination of direct application, such as in aquaculture and fruit production, as well as indirect release through sewage and animal manure. Antibiotic concentrations in many sewage-impacted rivers are predicted to be sufficient to drive antimicrobial resistance selection. Here we examine the potential for macrolide and fluoroquinolone prescribing in England to drive resistance selection in the River Thames catchment, England. We show that 63% and 73% of the length of the modelled catchment is chronically exposed to putative resistance-selecting concentrations of macrolides and fluoroquinolones, respectively. Our results reveal that macrolide and fluoroquinolone prescribing would need to decline by 77% and 85%, respectively, to protect freshwaters from resistance selection. We anticipate our model will provide a starting point for a more sophisticated national-scale assessment of the relationship between antibiotic prescribing and resistance selection in the environment. Improved antibiotic stewardship, alone, is unlikely to alleviate the identified challenge. Action is needed to substantially reduce antibiotic prescribing alongside innovation in sewage-treatment to reduce the discharge of antibiotics and resistance genes. Greater confidence is needed in current risk-based targets for antibiotics, particularly in mixtures, to better inform environmental risk assessments and mitigation.

## Introduction

The drive to reduce antibiotic use and misuse in human medicine had gained considerable traction since 2013 when Professor Dame Sally Davies, the Chief Medical Officer (CMO) for England and Chief Medical Advisor to the UK government published her Annual Report [1]. In Volume 2 of this report, she highlighted the “challenges and opportunities facing us in the prevention, diagnosis and management of infectious diseases”, which included the move to reduce inappropriate prescribing of antimicrobials. To this end, the National Health Service of England (NHS) proposed targeted reductions in antibiotic prescriptions as part of the Quality Premium Programme (QPP) [2].

The QPP in 2015/16 aimed to reduce antibiotic over-use and inappropriate prescribing through a reduction in:

1. the number of antibiotics prescribed in primary care by ≥1% from each Clinical Commissioning Group (CCG’s) 2013/14 value;
2. the proportion of broad-spectrum antibiotics prescribed in primary care. Specifically, the QPP aims to reduce prescriptions of co-amoxiclav, cephalosporins and fluoroquinolones by 10% (from each CCG’s 2013/14 value) as a percentage of the total number of antibiotics prescribed in primary care, or to be below the 2013/14 median proportion for English CCGs (11.3%), whichever represents the smallest reduction.

The NHS of England successfully reduced their antibiotic prescriptions in 2015/16 by 7.3% (37.03 million items to 34.34 million) as compared to 2014/15. The NHS also saw a reduction of 16% in broad-spectrum antibiotics (3.94 million items to 3.3 million). The goal for 2016/17 was, in part, to reduce total antibiotic prescribing in primary care by 4% (based on 2013/14) and broad-spectrum antibiotics by 20% (based on 2014/15). The majority of the reductions seen in 2016/17 were for amoxicillin, co-amoxiclav, and some cephalosporins, not macrolides or fluoroquinolones. The only exceptions being erythromycin, which saw a reduction in consumption in the primary care setting, of 0.078 DDDs and azithromycin which saw an increase of 0.023 DDDs per 1000 inhabitants per day in England [3].

Any reduction in antibiotic prescribing would result in a proportional reduction in antibiotics released into wastewater. This is because a significant proportion of antibiotics are conserved as they pass through the body before being excreted as a mixture of the parent compound and metabolites in the urine and faeces [4]. The gut bacteria from hundreds of thousands of NHS patients would have been enriched in bacteria harbouring antibiotic resistance, a phenomenon that is unavoidable upon consumption of antibiotics [5]. It is this chronic release of antibiotic and antibiotic-resistant bacteria from sewage effluent into freshwater that is thought to increase the prevalence of resistance genes in the environment substantially. This phenomenon is most evident downstream a sewage treatment plant (STP) discharge point as compared to upstream [6–11]. Recent evidence shows that even trace levels of sewage effluent can elevate the prevalence of markers of antibiotic resistance, i.e., class 1 integrons [12].

This study focused on two questions that explore the link between antibiotic use and environmental impact:

1. To what extent might current macrolide and fluoroquinolone prescribing drive antibiotic resistance selection in sewage-impacted rivers in southern England?
2. How much of a reduction in macrolide and fluoroquinolone prescribing might be required to alleviate the hazard of antibiotic resistance selection in rivers?

The study focused on macrolides and fluoroquinolones because: 1) they are two of the more persistent classes of antibiotics *in vivo* (Table 1) and the environment, and as such, are found in nearly all antibiotic surveillance studies [13–18]; 2) they have significant clinical relevance [19]; and, 3) macrolides are on the EU Watch List of Decision 2015/495/EU [20,21].

**Table 1.** Human excretion of fluoroquinolones and macrolides as a percentage of the parent compound.

| Class of Antibiotic | Antibiotics | Excretion (% of parent compound) |
| --- | --- | --- |
| Fluoroquinolones | Ciprofloxacin | 53.8 [22], 40 [23], 50 [24], 45-60 [25] |
|  | Levofloxacin | 71 (60-80) [22], 70 [23] |
|  | Moxifloxacin | 60 [26] |
|  | Norfloxacin | 61.5 [22], 30 [23] |
| Macrolides | Ofloxacin | 75.8 (60-80) [22], 100 [24] |
|  | Azithromycin | 50 [24], 50 [25], 8 [23] |
|  | Clarithromycin | 33.7 (14.4-60) [22], 20 [23], 0.18 [24] |
|  | Erythromycin | 35 (3.5-98) [22], 8 [23], 12-15 [25] |

The River Thames catchment (i.e., Thames Basin) in southern England was selected for this study as it is the most highly populous catchment in the United Kingdom (nearly 4 million people). It also might be seen as a realistic worst-case scenario, as on average, this part of England is in the 25^th^ quartile of predicted values of annual dilution factors, i.e., between 2.24 and 6.26 [27]. Many river stretches within the catchment offer little opportunity for dilution, maximising the impact of antibiotics found in the discharged sewage. Furthermore, the catchment has previously been the subject of several studies investigating predicted environmental concentrations of pharmaceuticals [17,28–30].

## Materials and methods

### Prescription data

Macrolide prescribing data for each month of 2015/16 were acquired from the NHS Business Service Authority (NHSBSA), i.e., azithromycin, clarithromycin and erythromycin (Fig 1a). Fluoroquinolone prescribing data for each month of 2015/16 were also acquired from the NHSBSA, i.e., ciprofloxacin, levofloxacin, moxifloxacin, norfloxacin and ofloxacin (Fig 1b). Prescription data were acquired from four Clinical Commissioning Groups (CCGs): Oxfordshire, Gloucestershire, Swindon and Wiltshire, all of which are situated in the upper River Thames catchment (Fig 2). It was not possible to reliably assign CCG prescriptions to STPs in the lower Thames catchment owing to the small geographic size of the CCGs and high population density. As such, all other CCGs within the catchment were assigned the prescription rate of NHS Oxfordshire CCG.

**Fig 1.**
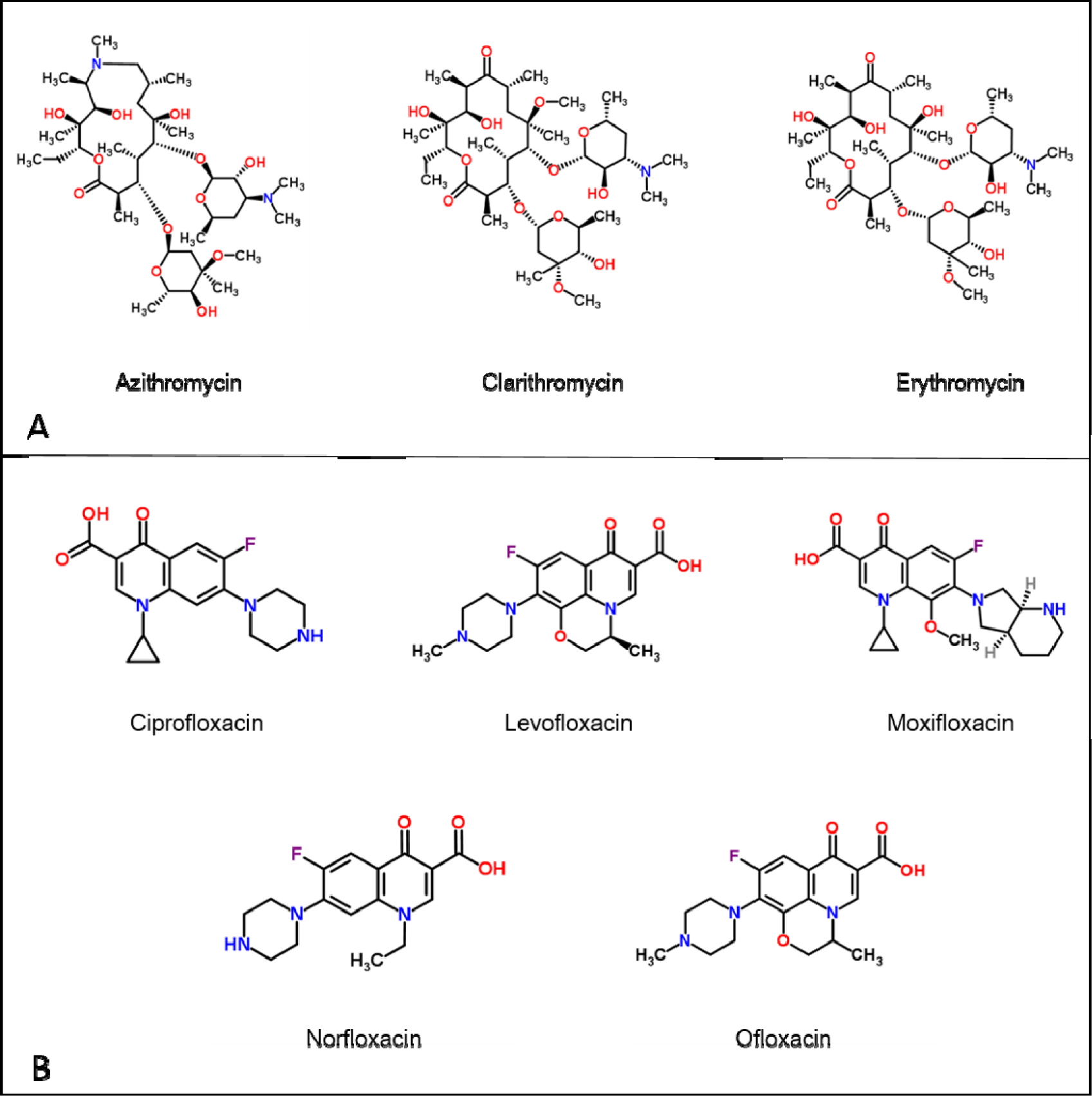
Macrolides (A) and fluoroquinolones (B) included in the study.

**Fig 2.**
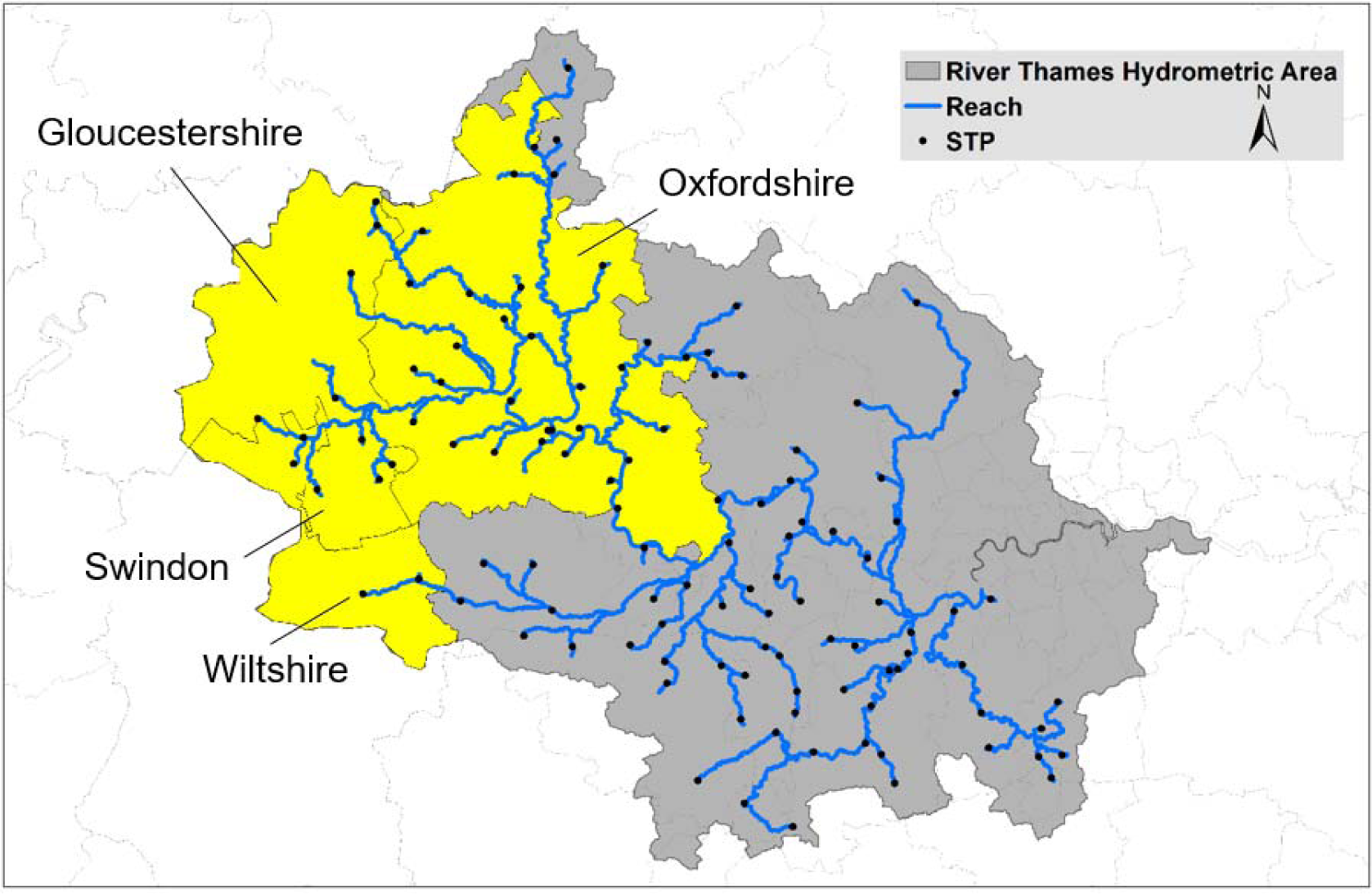
Hydrometric area of Thames Region with the four CCGs in the upper Thames Catchment denoted in yellow.

NHS prescription data for each CCG per month in 2014/15 were converted to units of kg (Table 2), and then moles (Table 3) of macrolide and fluoroquinolone (Table 2) and subsequently normalised to the population within each CCG (Table 4). In the case of erythromycin, the three forms of erythromycin (erythromycin, erythromycin ethyl succinate and erythromycin stearate) were converted individually and summed.

**Table 2.** Mass of macrolides and fluoroquinolones prescribed in the study CCGs in 2015/16.

|  |  | Oxfordshire | Gloucestershire | Swindon | Wiltshire |
| --- | --- | --- | --- | --- | --- |
|  |  | (kg) | (kg) | (kg) | (kg) |
| Macrolides | Azithromycin | 21.8 | 18.9 | 4.5 | 12.6 |
|  | Clarithromycin | 179.5 | 322.7 | 66.7 | 198.0 |
|  | Erythromycin | 97.3 | 80.7 | 50.6 | 82.3 |
|  | Erythromycin ethyl succinate | 120.8 | 79.0 | 32.3 | 34.4 |
|  | Erythromycin stearate | 12.4 | 5.4 | 4.3 | 5.0 |
|  | Total of macrolides (kg) | 431.8 | 506.7 | 158.4 | 332.3 |
| Fluoroquinolones | Ciprofloxacin | 93.8 | 94.6 | 29.5 | 72.2 |
|  | Levofloxacin | 0.3 | 4.6 | 0 | 0.3 |
|  | Moxifloxacin | 0.5 | 0.1 | 0.1 | 0.2 |
|  | Norfloxacin | 0 | 0 | 0.1 | 0 |
|  | Ofloxacin | 2.2 | 4.1 | 0.7 | 1.0 |
|  | Total of fluoroquinolones (kg) | 96.8 | 103.4 | 30.4 | 73.7 |

**Table 3.**
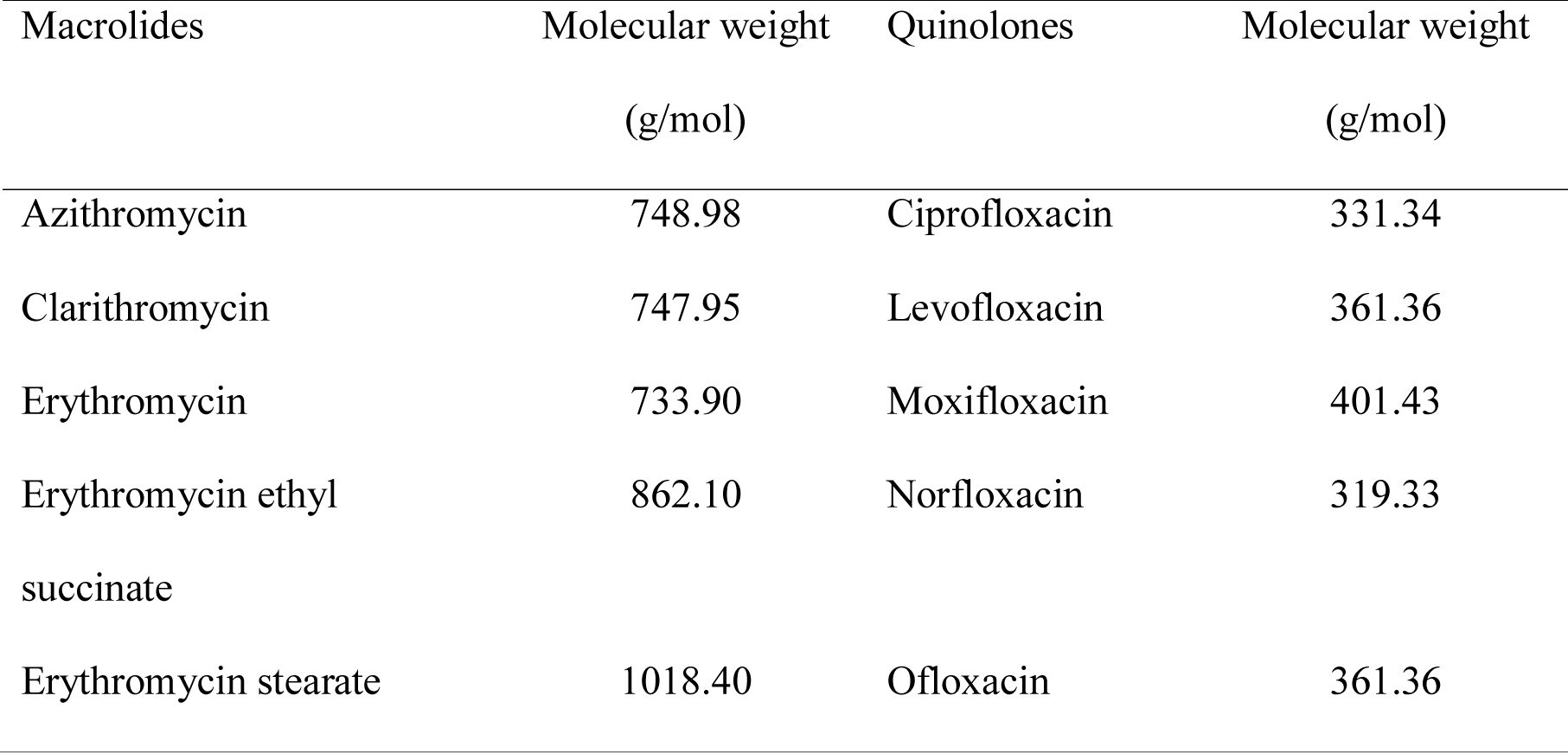
Molecular weights of macrolides and fluoroquinolones.

**Table 4.**
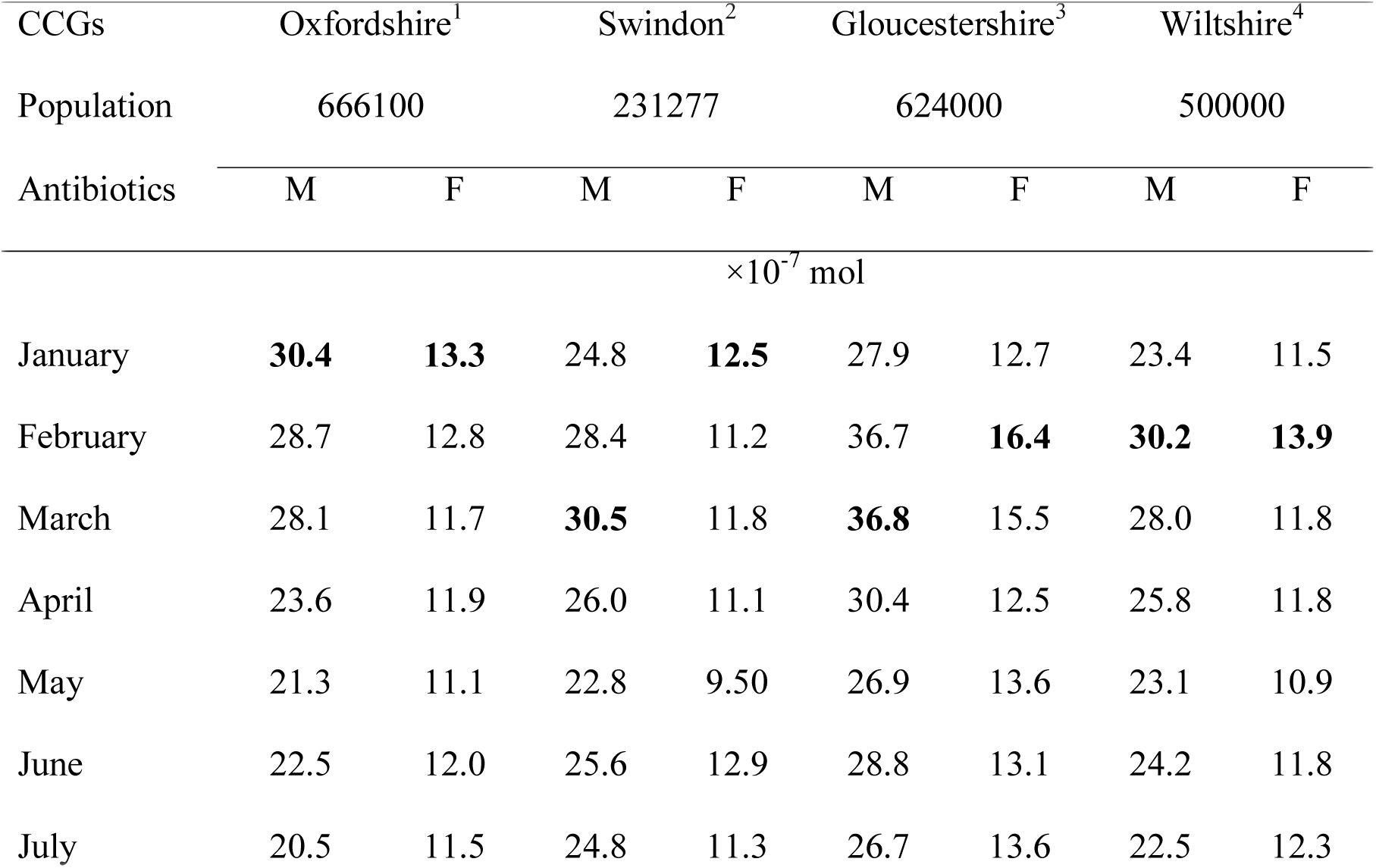
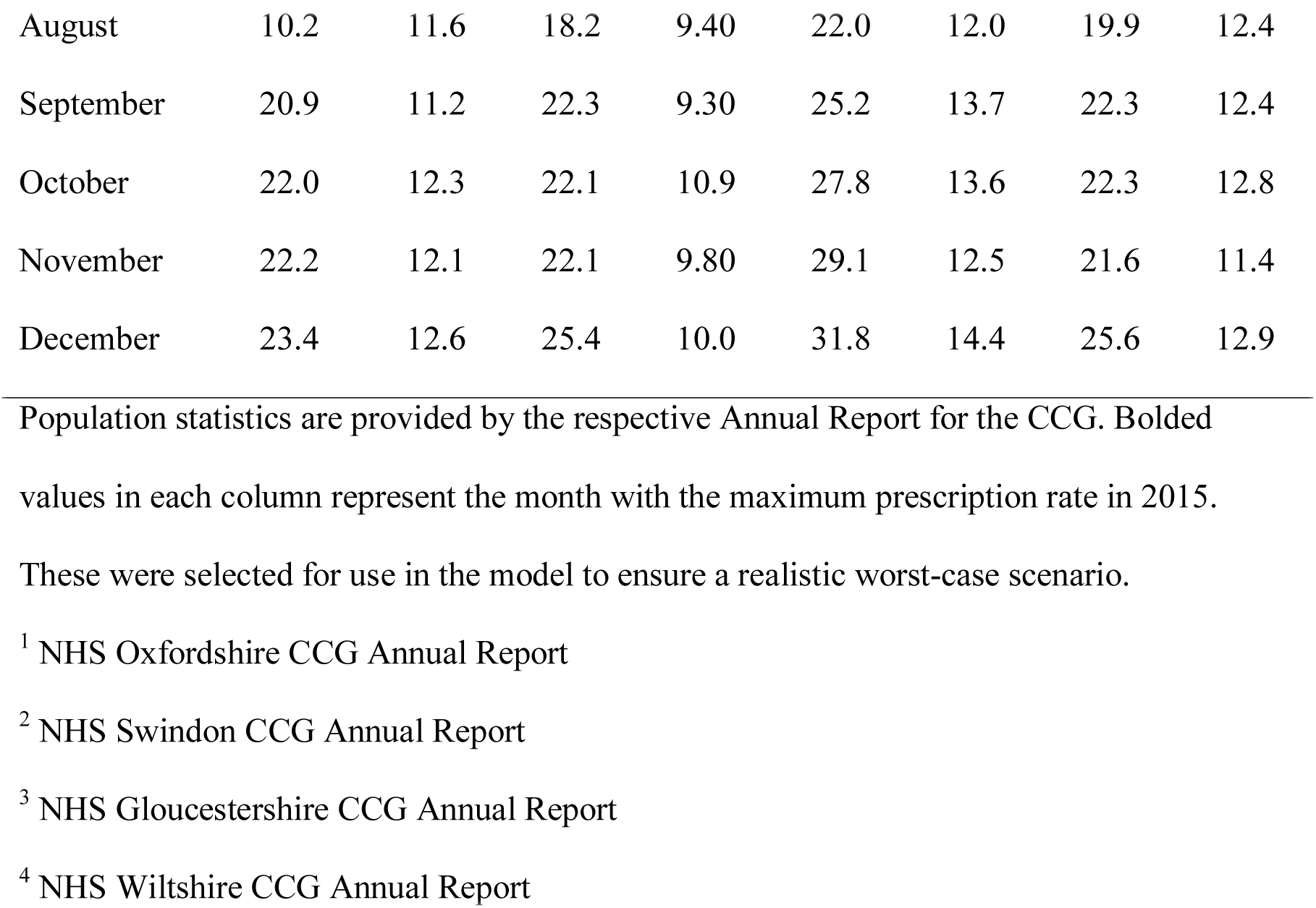
Moles of macrolides (M) and fluoroquinolones (F) prescribed capita^−1^ day^−1^ month^−1^ within each CCG.

### Antibiotic excretion rate

The excretion of antibiotics in the faeces and urine depends on its metabolism *in vivo*, which is a function of health, age, gender, ethnicity, as well as dosage and mode of administration, e.g., capsule, injection. Given the large uncertainty associated with estimating excretion rates across diverse populations, an average rate acquired from the literature was used: macrolide excretion rate of 32.2% and 64.2% for fluoroquinolones (Table 1).

### Antibiotic loss in STPs

The antibiotic loss in STPs was modelled using STPWIN model within the Estimation Program Interface (EPI) Suite^TM^ 4.0, using the Biowin/EPA draft method for determining half-life data as previously described [29]. Estimates of loss in STPs was also acquired from the literature [31–36], with some relevant data acquired from STPs within the River Thames catchment found in Table 5. These data support the use of 50% loss during STP passage, which is a compromise between measured loss and the high variability reported within the literature.

**Table 5.** Measured antibiotics concentration in STPs and rivers within the Thames Catchment (ng/L).

|  | Macrolide |  |  | Fluoroquinolone |  |  |
| --- | --- | --- | --- | --- | --- | --- |
|  | CLAR | ERY | AZO | CIP | NOR | OFL |
| Eff – Oxford* | 98 (338 – 1504) |  | 156 (69 – 264) |  |  |  |
| Eff – Didcot* | 104 (160 – 305) |  | 110 (74 – 216) |  |  |  |
| Eff – Cholsey* | 181 (128 – 321) |  | 91 (48 – 135) |  |  |  |
| Eff – Benson* | 152 (87 – 254) |  | 76 (35 – 133) |  |  |  |
| Eff – Benson** | 50 | 244 | 34 | 14 | 25 | 195 |
| Eff – Oxford** | 91 | 236 | 30 | 52 | 21 | 23 |
| Thames var.*** | 292, 30 | 448, 58 | 51, 21 | 46, 20 | 45, 9 | 17, 11 |

Eff sewage effluent, CLAR clarithromycin, ERY erythromycin, AZO azithromycin, CIP ciprofloxacin, NOR norfloxacin, OFL ofloxacin
\* Mean (highest – lowest) measured [37]
\*\*24-h mean concentration [17]
\*\*\*Max and mean concentration for 21 locations at 7 time points [17]

### Modelled environmental concentrations of antibiotics

Low Flows 2000 Water Quality eXtension (LF2000-WQX) model [30,38] is an extension to the LF2000 [39]. LF2000 is a decision support tool designed to estimate river flow at gauged and ungauged sites and to assist regional water resources and catchment management. The LF2000-WQX software is a geographical information-based system that combines hydrological models with a range of water-quality models, including a catchment-scale water-quality model. This model generates spatially explicit statistical distributions of down-the-drain chemicals for both conservative and degradable compounds. It uses a Monte Carlo model approach to combine statistical estimates of chemical loads at specific emission points (e.g., STP) with estimated river flow distributions for the whole river network of interconnected model reaches (a reach is the river stretch between model features, e.g., major tributaries, STPs). The hydrometric area of the River Thames catchment and the STPs included in LF2000-WQX (black points) can be seen in Fig 2. Thus, working from the upstream reaches at the head of the river network to the outlet of the river basin, the model accounts for the accumulation of point antibiotic loads and the water in which these loads are diluted. Degradable chemicals might be removed from STPs and the river water. The latter is represented by a non-specific dissipation process assuming first-order kinetics., Antibiotics were assumed to persist in the river for at least one day, thereby providing a realistic worst-case scenario.

In summary, LF2000-WQX was parameterised with the highest monthly fluoroquinolone and macrolide prescription rate for each of the four CCGs (see bolded rates in Table 4). Macrolide and fluoroquinolone loss was accounted for upon excretion (32.2% and 64.2%, respectively) and before discharge from STPs into the receiving river (50%). As such, the mass of macrolide and fluoroquinolone consumed was reduced by 83.9% and 67.9%, respectively, upon discharge into the adjacent river. The antibiotic load was subsequently diluted within the river, parameterised with the mean annual river flows within LF2000-WQX.

### Risk-based targets for antibiotics

Risk-based management targets for antibiotics in freshwater would ideally be set at concentrations that are below the lowest concentration that allows antibiotics to select for antibiotic resistance genes. Such effect concentrations have been determined experimentally in a limited number of lab-based studies, operationally termed minimum selection concentrations (MSC). MSCs typically range between 0.1 to 10 µg of antibiotic/L [40–52].

Modelled predicted no-effect concentrations (PNECs) were acquired from Bengtsson-Palme and Larsson (2016) owing to the absence of empirically-determined MSCs for most of the study antibiotics. The PNECs derived in Bengtsson-Palme and Larsson (2016) were selected to inform antibiotic discharge limits from pharmaceutical manufacturing by the AMR Industry Alliance [53]. The PNECs represent the threshold concentration of an antibiotic, above which, there is a heightened hazard of antibiotic resistance selection. PNECs were derived from the European Committee on Antimicrobial Susceptibility Testing (EUCAST) database of minimum inhibitory concentrations to form species sensitivity distributions [29]. The authors selected the concentration of each antibiotic representing the 1% potentially affected fraction (PAF). A safety factor of 10 was added to this 1% PAF to account for the observation that experimentally-derived resistance selection thresholds tend to be approximately an order of magnitude lower than the MIC, while also offering an added level of protection to the estimate. The 111 antibiotic thresholds reported in Bengtsson-Palme and Larsson 2016 ranged from 0.008 µg/L to 64 µg/L.

PNECs for fluoroquinolones ranged from 0.064 to 0.5 µg/L and for macrolides 0.25 to 1 µg/L (Table 6). When modelled concentrations of macrolides and fluoroquinolones exceeded the PNEC for the most ‘potent’ antibiotic within each class, e.g., ciprofloxacin for fluoroquinolones and azithromycin/clarithromycin for macrolides, the stretch of river was denoted as being ‘at risk’ for resistance selection. When the modelled concentrations exceeded the PNEC for the least ‘potent’ antibiotic within each class, e.g., norfloxacin/ofloxacin for fluoroquinolones and erythromycin for macrolides, the stretch of river was denoted as being at a ‘critical risk’ of resistance selection.

**Table 6.** Modelled PNECs for macrolides and fluoroquinolones [**40**].

| Antibiotic class | Antibiotic | PNEC<br>($\mu\text{g/L}$) | PNEC<br>($10^{-10}$<br>mol/L) |
| --- | --- | --- | --- |
| Fluoroquinolones | Ciprofloxacin | 0.064 | 1.9 |
|  | Levofloxacin | 0.25 | 6.9 |
|  | Moxifloxacin | 0.125 | 3.1 |
|  | Norfloxacin | 0.5 | 15.7 |
|  | Ofloxacin | 0.5 | 13.8 |
| Macrolides | Azithromycin | 0.25 | 3.3 |
|  | Clarithromycin | 0.25 | 3.3 |
|  | Erythromycin | 1 | 9.8 |

## Results

This study aimed to understand the degree of resistance selection that might be occurring in the River Thames catchment under current macrolide and fluoroquinolone prescribing practice by the NHS as well as after foreseeable and aspirational reductions in prescribing.

The objective was to estimate the level of reduction needed in NHS antibiotic prescribing that might be necessary to substantially reduce the macrolide and fluoroquinolone-resistance selection burden created in the river environment.

Predicted environmental concentration (PEC) for macrolides and fluoroquinolones prescribed in 2015/16 revealed a maximum concentration of 4.3 µg/L and 2.0 µg/L, respectively. Hazard characterisation revealed 279 out of 457 reaches in the catchment are ‘at risk’ or ‘critical’ for macrolide resistance selection (Fig 3a)—equating to 1155 of 1398 km of the modelled River Thames catchment (63.2% of the total catchment length modelled). Similarly, 311 reaches out of 457 were ‘at risk’ or ‘critical’ for fluoroquinolone resistance selection (Fig 3b)— equating to 1026 of 1398 km (73.4% of the modelled river length in the catchment).

**Fig 3.**
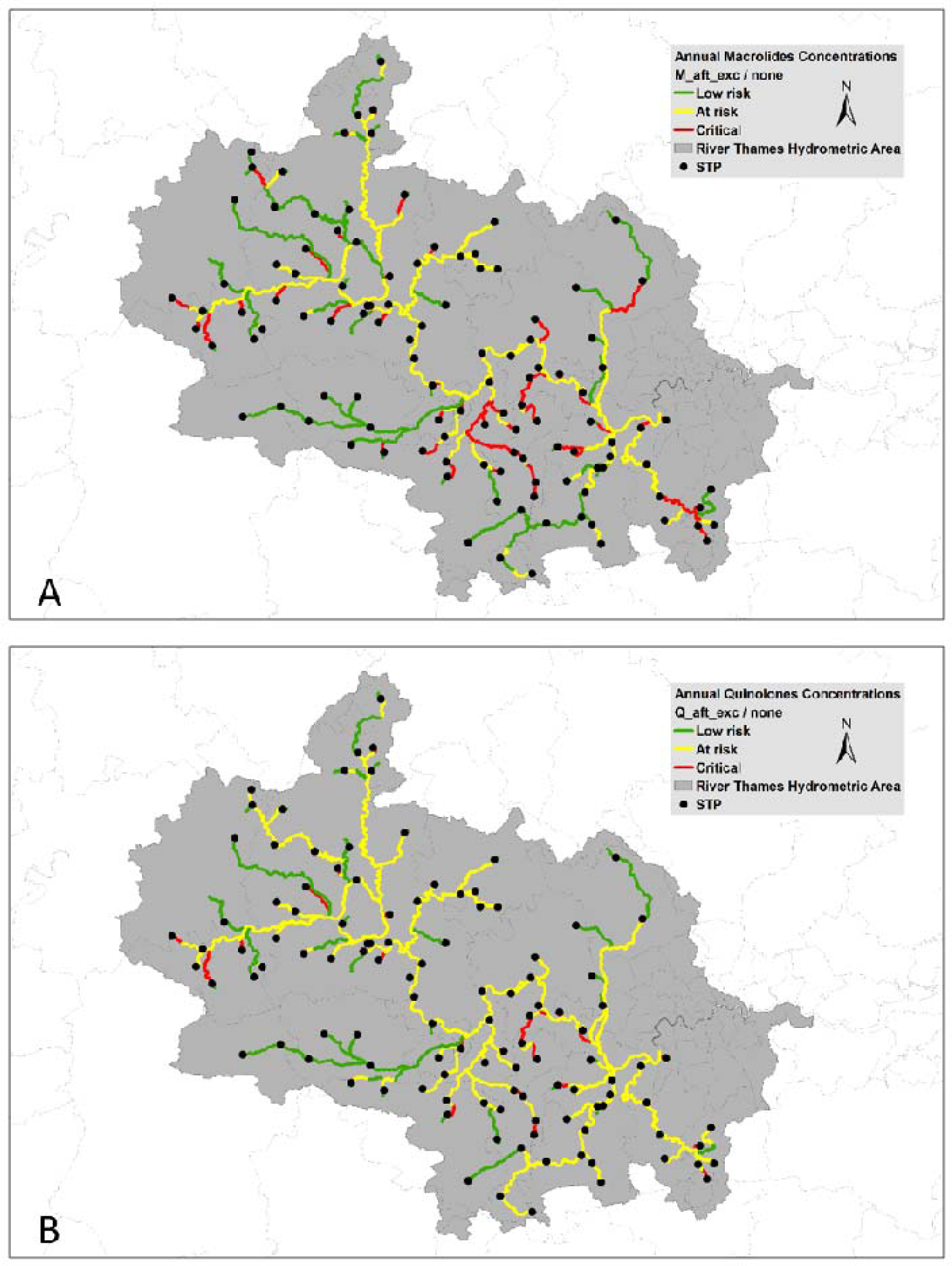
Resistance selection risk characterisation for macrolides (A) and fluoroquinolones (B) using 2015/16 prescription statistics.

Consistent with the 2015/16 QPP goals, a 4% reduction in macrolide prescribing saw a 3.8% reduction in the length of the modelled Thames catchment ‘at risk’ for macrolide resistance selection (273 reaches and 830 km, equal to 59.4% of the catchment), and a 0.1% reduction in the length of the modelled Thames catchment ‘at risk’ for fluoroquinolone resistance selection (310 reaches and 1025.1 km, equal to 73.3% of the catchment).

A more ambitious 20% reduction of the 2015/16 prescription rate of macrolides resulted in an 8.9% reduction in the length of the modelled Thames catchment ‘at risk’ for macrolide resistance selection (254 reaches and 759 km, equal to 54.3% of the catchment). A 20% reduction in fluoroquinolones resulted in a 5.4% reduction in the length of the modelled Thames catchment ‘at risk’ for fluoroquinolone resistance selection (296 reaches and 949 km, equal to 68.0% of the catchment).

A sensitivity analysis was conducted to identify the NHS prescribing rate that protects ≥90% of the length of the modelled Thames catchment from macrolide and fluoroquinolone resistance selection. A target of 90% was a pragmatic decision, as many rivers in England are highly impacted by, or solely composed of, sewage effluent—particularly in headwaters. Hence, a target of 100% would be unrealistic and necessitate even more dramatic reductions in antibiotic use or zero effluent discharge.

The sensitivity analysis revealed that macrolide prescriptions must decline by 77% to protect 90.4% of the length of the Thames catchment from macrolide resistance selection (Table 6; Fig 4a). Even at this much-reduced prescription rate, there were still 13 reaches with a total length of 8.5 km (0.6%) remaining ‘critical’ for macrolide resistance selection (Fig 4a). Fluoroquinolone prescriptions would need to be reduced by 85% to alleviate the selection risk in 90% of the catchment (Table 6; Fig 4b).

**Fig 4.**
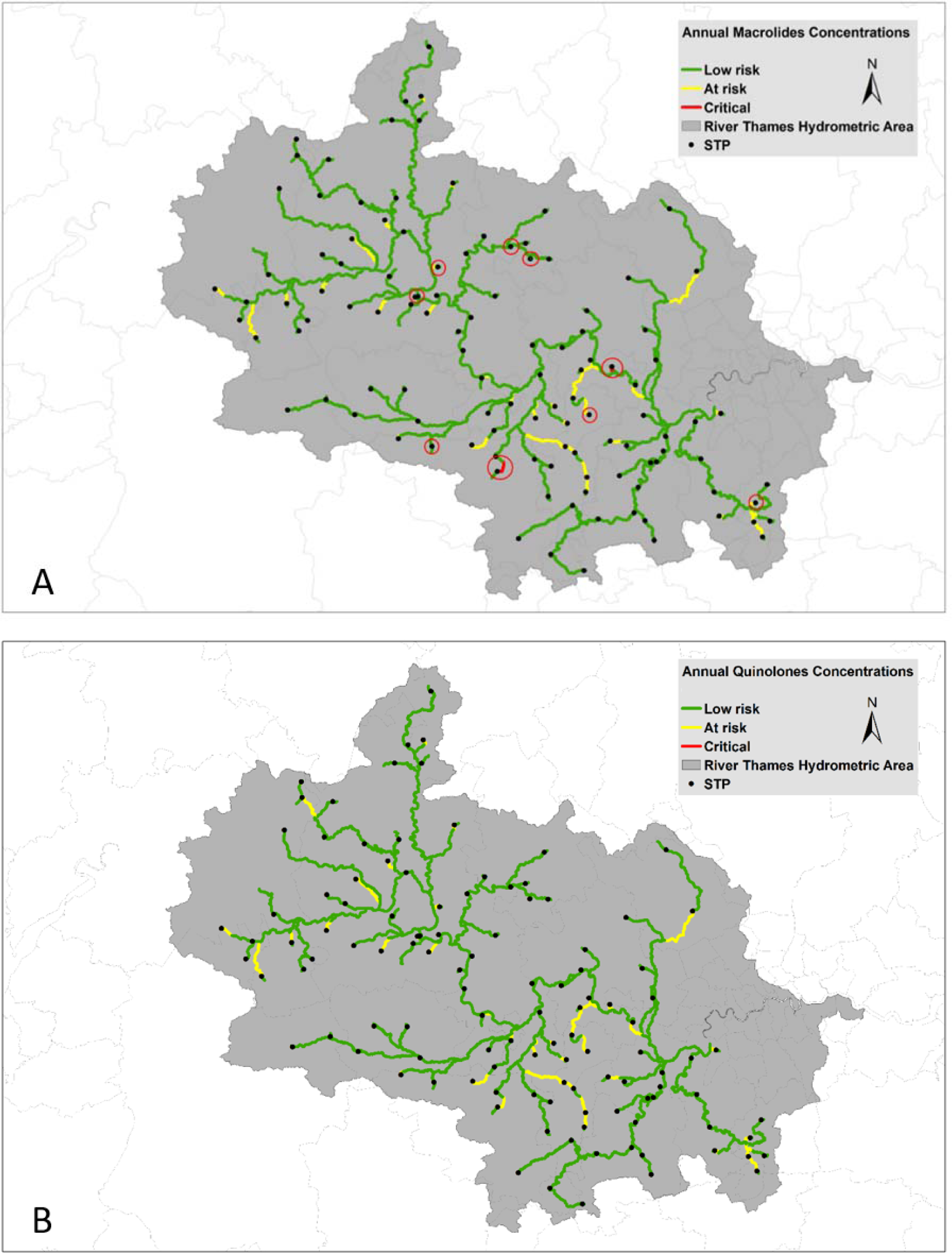
Hazard characterisation for macrolide (A) and fluoroquinolone (B) resistance selection after a reduction of 77% and 85% in prescriptions, respectively, on 2015/16 rates. Red circles in (A) indicate nine locations where the concentrations remained ‘at critical’ levels.

## Discussion

The goal of this study was to evaluate how antibiotic prescribing, alone, contributes to antibiotic resistance selection in the aquatic environment. Although such a perspective is fundamentally naïve given the omission of antibiotic use in animals and other pollutants that can co-select for antibiotic resistance, it provides a conservative estimate of our current state of the environment and a starting point for discussing mitigation [54–56].

A non-trivial proportion of the resistance genes found downstream of STPs originate from human gut bacteria, many of which have acquired resistance *in vivo*, perhaps during antibiotic treatment of human infections. Antibiotic-resistance genes disseminated from anthropogenic sources into the environment are pollutants [57,58] maintained and spread through: 1) survival of enteric bacteria in the environment and wildlife [59–61]; 2) horizontal gene transfer of resistance genes from enterics into indigenous environmental bacteria [62]; and 3) selection and co-selection taking place *in situ* [63,64].

Evidence of these processes occurring in the River Thames catchment has been reported in the literature. Extended-spectrum β-lactamase-producing *Escherichia coli* (bla_CTX-M-14_) belonging to the clinically important O25b:H4-ST131 lineage was isolated from water from the River Thames in west London [65]. Amos *et al.* 2015, analysed sediment samples from 13 sites across the River Thames basin for class 1 integron prevalence and cefotaxime- (third-generation cephalosporin)-resistant *E. coli* [6]. The authors reported antibiotic resistance was strongly correlated to the proximity and size of upstream STPs. Lehmann *et al.* 2016, showed a rapid increase in class 1 integrons in natural river biofilms, within the Thames catchment, exposed to trace levels of additional STP effluent [12]. Hence, the presence of elevated levels of antibiotic resistance in sewage-impacted rivers is most likelya function of DNA pollution nested over a background of resistance selection driven by antibiotic and chemical pollution.

### Strengths and Limitations

To our knowledge, this study is the first attempt to model the impact of current and aspirational antibiotic prescribing in England on antibiotic resistance selection in freshwater rivers. The study’s approach reflects the state of our knowledge at this time, and as such, is limited in many ways.

### Heterogeneity of population

The results do not reflect the granular differences in prescribing that are seen between small CCGs as there would be no way to attribute an antibiotic user to a particular STP confidently. It is expected that if such an effort were possible, it would highlight elevated antibiotics in STPs that have: older demographics (e.g., care homes [66]), hospitals [67,68], and areas of inhabitants with elevated international travel [69,70]. Should these STPs be located at or near headwaters, it would likely result in elevated hazards for antibiotic resistance selection resulting from lower effluent dilution. Conversely, when these high-antibiotic users are located in stretches where there is sufficient dilution, the selection hazard would be alleviated, somewhat.

### Varied antibiotic stewardship within antibiotic class

In this model, we applied any reductions in antibiotic prescribing, i.e., 4 or 20%, uniformly across all antibiotics within the class. However, use of antibiotics within a class are unlikely to be uniformly reduced. As a result, some reductions in prescribing could delay or hasten the time to achieve ‘low risk’. For example, ciprofloxacin is the most ‘potent’ fluoroquinolone in its class, with a PNEC of 64 ng/L; ofloxacin and norfloxacin are less ‘potent’, with a PNEC of 500 ng/L. In other words, ciprofloxacin is 7.8125 times more potent than ofloxacin and norfloxacin. Hence, the benefit gained from reducing 1 unit of ciprofloxacin necessitates a reduction in 7.8125 units of ofloxacin or norfloxacin. As ciprofloxacin represents the majority of fluoroquinolone prescribed (96.9%; Table 2), any reduction in ciprofloxacin is likely to have the maximum impact on fluoroquinolone resistance selection. Similarly, macrolides azithromycin and clarithromycin have PNECs of 250 ng/L, four times more potent than erythromycin (1000 ng/L). Prescription of macrolides varies considerably between the four study CCGs (Tables 2 and 4), with clarithromycin (Gloucestershire and Wiltshire CCG) or erythromycin (Oxfordshire and Swindon) being the most frequently prescribed macrolide. Hence, the CCGs that are overprescribing clarithromycin will find reduced macrolide prescriptions potentially more impactful owing to the higher potency of clarithromycin relative to erythromycin.

### Reliability of PNEC

The PNECs employed in this study have recently emerged as modelled estimates for assessing the hazard of resistance selection [40]. To date, modelled PNECs are consistent with the growing literature base [40–52], and as such, have been adopted by the AMR Industry Alliance for establishing regulatory thresholds in antibiotic manufacturing waste effluent [53]. There is a risk that the modelled PNECs are too protective, and would encourage potentially costly and unnecessary mitigation. Conversely, modelled estimates of single antibiotics might also underestimate selection risk, as antibiotic (pollutant) mixtures can act synergistically, as discussed below. The academic community will need to continually challenge the suitability of these PNECs and ensure they are fit for purpose.

#### Mixture effects

Recent experimental evidence suggests antibiotics can act synergistically and antagonistically when in mixtures [71,72]. Tekin *et al.* 2018, reported that synergistic and antagonistic interactions increased in frequency with the number of drugs in the bacteria’s environment [72]. Hence, a non-trivial number of the modelled PNECs used in this study could be over- as well as under-estimating the threshold for resistance selection when applied to sewage-impacted rivers.

Brochado *et al.* 2018, demonstrated that many synergies were found within the same class of antibiotic, in particular, where the same cellular process was targeted. The authors postulated that by targeting the same functional process at different steps, drug combinations of the same class could bypass the apparent redundancy of having multiple antibiotics within the same class and exhibit a synergistic effect [71]. The authors also demonstrated that many of the drug antagonisms involve antibiotic resistance mechanisms that modulate intracellular drug concentrations and not direct interactions of the primary drug targets. Hence, some antibiotic combinations were less inhibitory than expected (i.e., non-additive) because the cell’s response to one drug helped to buffer the effects from the second drug. For example, a decrease in the intracellular concentration of one drug by an efflux pump, can potentially also decrease the intracellular concentration of the interacting drug [71]. Notably, the authors found that the drug-drug interactions were often conserved within species (70%), but with 13–32% of the interactions strain-specific, and poor conservation across species (5% of drug-drug interactions). It is premature to apply any specific drug-drug interactions to this model, but future efforts should attempt to account for mixture effects as part of a holistic hazard assessment.

### Mixture Effects with Non-Antibiotics

Another limitation of this study is that it does not take into account the presence of many potentially relevant chemicals that are known to be present in sewage and sewage-impacted rivers, such as:

1. antibiotics from agriculture and animal use [73];
2. biocides, which have been shown to co-select for antibiotic resistance genes [74–76];
3. metals, which have been shown to co-select for antibiotic resistance genes [77–80];
4. other classes of pharmaceuticals that might have synergistic or antagonistic impacts on resistance selection [71,81,82], and
5. herbicides and pesticides that have been shown to co-select for resistance genes [83,84].

It is impractical to experimentally test all the chemicals found in sewage in isolation and mixtures for the threshold concentration that allows for resistance selection, as there are simply too many. However, our environment can be assayed for such insight, by measuring the diversity and quantity of antibiotic resistance genes in the river environment and their relationship to STPs, farms, hospitals, aquaculture, urban runoff, etc. Where ‘pristine’ environments still exist, they might serve as benchmarks for qualifying and quantifying ‘natural’ resistance gene prevalence and biogeography. However, this might prove difficult as wildlife have been implicated in the dissemination of human clinical resistance genes, making ‘pristine’ locations potentially unattainable [60]. Quantifying ‘natural’ resistance gene abundance and prevalence remains critical for hazard assessment and establishing mitigation targets.

### LF2000-WQX Limitations

LF2000-WQX generates PECs from STPs characterised by dry weather flows of >5000 m^3^/d. A majority of the small STPs not included in the model are located in the upper-most reaches of the catchment where flows are typically very low. Headwaters impacted by small STPs are likely to have a disproportionate impact on seeding the catchment with pharmaceuticals as well as antibiotic-resistant bacteria, the former of which would further add to the total load of macrolides and fluoroquinolones in the catchment. The omission of these STPs is unlikely to increase the antibiotic load in the catchment substantially; however, it might impact the DNA ‘seeding’ and maintenance of antibiotic resistance.

### Aspirational Reductions in Prescribing

In this study, we projected a decline in antibiotic prescribing without considering the likely change in other important factors that would occur concurrently with a gradual reduction in prescribing. An increasing and ageing UK population will consume more pharmaceuticals per capita increasing the antibiotic load. Dryer summers, resulting from climate change, might impact the frequency PNECs are exceeded owing to lower dilution [85,86]. More intense storms predicted in a changing climate would lead to more frequent and longer combined sewage overflows (CSOs), which deposit raw sewage into the adjacent river, thereby elevating the release of resistant bacteria, genes and chemical pollutants into the environment. CSOs, septic tanks, and farm runoff represent potentially significant sources of antibiotics and resistance genes which would benefit from being included in future models. However, such data is not currently available, representing a large knowledge gap for risk assessment.

#### Validation of PECs

Previous research using LF2000-WQX to generate PECs, e.g., steroid estrogens [30,87], glucocorticoids [88], cytotoxic chemotherapy drugs [89] and antivirals [29], provides reassurance that the river network representation within LF2000-WQX is accurate (i.e., network, flows, dilution and STPs). The modelled antibiotic concentrations from LF2000-WQX, denoted by the PNEC thresholds in Table 6, are within the same range as measured environmental concentrations detailed in Table 7, lending credibility to the model outputs.

**Table 7.** Sensitivity analysis for level of reduction in prescriptions needed to protect 90+% of the length of the modelled River Thames catchment from resistance gene selection.

|  | % Reduction in Prescriptions | Number of reaches 'at risk' or 'critical' | Length 'at risk' or 'critical' (km) | % Length 'at risk' or 'critical' |
| --- | --- | --- | --- | --- |
| Macrolides | 80 | 69 | 110 | 7.9 |
|  | **77** | **82** | **134** | **9.6** |
|  | 76 | 86 | 157 | 11.2 |
|  | 75 | 93 | 169 | 12.1 |
| Fluoroquinolones | **85** | **82** | **135** | **9.6** |
|  | 84 | 88 | 157 | 11.2 |
|  | 83 | 96 | 175 | 12.5 |
|  | 80 | 118 | 215 | 15.4 |

#### Antibiotic Stewardship

Reduction of antibiotics in the environment can be achieved through a range of mitigating actions, the first of which can be through improved antibiotic stewardship. The results of this study indicate that current goals for antibiotic stewardship do not go far enough [90,91]. In Europe, the Netherlands has among the most restricted antibiotic stewardship, with nearly 50% fewer antibiotics prescribed in the community/primary care sector (10.06 DDD/1000 inhabitants/d) than in the UK (19.09), in 2017 (https://ecdc.europa.eu/en/antimicrobial-consumption/database/country-overview). Macrolide prescribing in the Netherlands (1.38 DDD/1000/d) was only 47% that of the UK (2.90 DDD/1000/d). The extent to which the Netherlands still overprescribes and misprescribes macrolides would be highly instructive in defining safe limits for further reductions in macrolide prescribing. Notably, the UK has a lower fluoroquinolone prescribing rate in the community (0.44 DDD/1000/d) as compared to the Netherlands (0.75). Hence, there are important lessons to be mutually shared across Europe and more widely on how to optimise antibiotic prescribing—a point not lost on the Advisory Committee on Antimicrobial Prescribing, Resistance and Healthcare Associated Infection [92] and Public Health England [93].

The UK has reported between 8.8% and 23.1% of all systemic antibiotic prescriptions in English primary care as inappropriate, with some high prescribing practices in England capable of reducing antibiotic prescriptions by as much as 52.9% [93]. In a linked study where the authors and an expert panel explored the ‘appropriateness’ of antibiotic prescription, the authors found that substantially higher proportions of patients received antibiotics than was deemed ‘appropriate’, with the respiratory ailments showing the largest contrast between actual and ideal: acute cough (41% vs 10%, respectively), bronchitis (82%:13%); sore throat (59%:13%); rhinosinusitis (88%:11%); and acute otitis media in 2- to 18-year-olds (92%:17%) [94]. Such a reduction in prescribing will be beneficial to reducing resistance selection in humans, but, as shown in this paper, might not go far enough (Table 8).

**Table 8.** Summary of modelled impact of antibiotic prescribing on antibiotic resistance gene selection in sewage-impacted freshwater.

| Scenarios | Macrolide | Fluoroquinolone |
| --- | --- | --- |
|  | % of modelled catchment length >PNEC |  |
| Antibiotic prescribing from 2015/16 | 63 | 73 |
| 4% Reduction on 2015/16 | 59 | 73 |
| 20% Reduction on 2015/16 | 54 | 68 |
|  | % reduction in prescribing required to achieve target |  |
| ≥90% of catchment <PNEC | 77 | 85 |

### Mitigation through Innovation and Investment in STPs

A step-change in the way we currently handle our wastewater in needed to tackle the challenge of antibiotic and DNA pollution. One option is to set emission limits on the concentration of antibiotics, in much the same way the pharmaceutical industry has voluntarily adopted for its manufacturing supply chain [53]. The setting of strict antibiotic emissions limits will have the desired effect of greatly reducing the environmental impact of antibiotics, with, arguably, several serendipitous implications for other chemical hazards. For example, any technological solution for the treatment of wastewater that is effective in removing a wide range of antibiotics will also likely reduce the load of estrogens and estrogen-mimicking chemicals which has been a growing environmental concern for several decades [95–97]. As such, it is entirely likely that the substantial cost associated with tackling estrogens and estrogen-mimicking chemicals [98] can be shared with the challenge of reducing AMR in the environment.

Moreover, there is a rapidly growing list of chemicals that are found within sewage effluent that has been shown to select or co-select for ARGs, i.e., antiepileptics [81], biocides/disinfectants [99,100], metals [101]. Hence, engineering solutions to antibiotic removal from wastewater can spread the cost across a very wide range of pollutants, thereby alleviating a substantial range of ecotoxicological hazards, in addition to resistance selection. Spreading the mitigation costs across a range of challenges might also be more politically and socially tractable, owing to the significant cost associated with improving our treatment of wastewater.

Although not immediately within the scope of this paper, it is relevant to highlight that any engineering solutions employed for the removal of antibiotics from sewage effluent might be ineffective in reducing the DNA pollution that ARGs represent. Co-development of engineering solutions to tackle the chemical and biological drivers of ARGs in the environment is required to reduce the environmental pressure caused by our wastewater. In addition, the by-products of STP, sludge, which contains antibiotic resistance, antibiotics and non-antibiotic chemical drivers of resistance are currently amended to land, representing additional risks to humans and the environment [54], but are out of the study’s scope.

## Conclusion

This study explores the reduction in macrolide and fluoroquinolone prescribing needed to alleviate the modelled hazard from antibiotic resistance selection in sewage-impacted rivers. It is unclear if the projected reductions in antibiotic pollution of 77 to 85% could be achieved solely through reduced prescribing by the NHS. Environmental targets could be more readily achieved by a holistic, integrated AMR action plan, which constrains and optimises antibiotic prescribing, while also addressing the chronic release of antimicrobials, biocides, metals and resistance genes from STP effluent.

## Acknowledgements

We gratefully acknowledge help from Elizabeth Beech from the National Health Service of England, and Richard Williams and Holly Tipper of the Centre for Ecology & Hydrology, United Kingdom.

